## Supplementary figures and images for "Spatial and temporal transcriptomics of SHH-medulloblastoma with chromothripsis identifies multiple genetic clones that resist to treatment and lead to relapse"

### Figure S1

Supplementary Figure 1

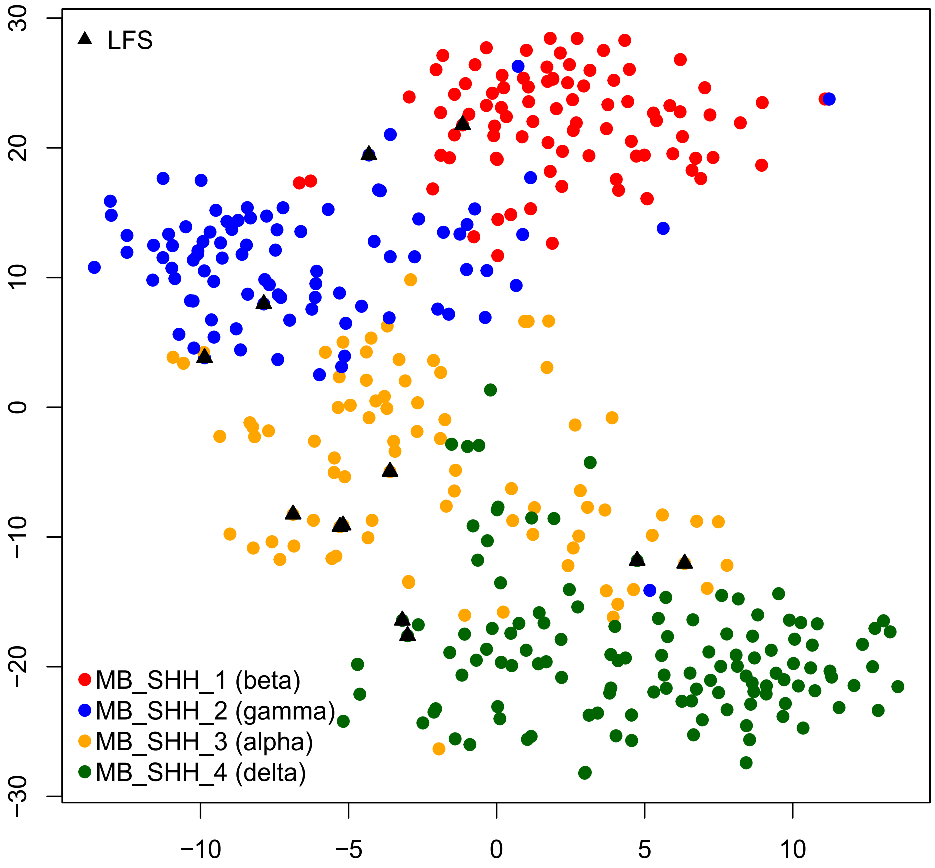

LFS1

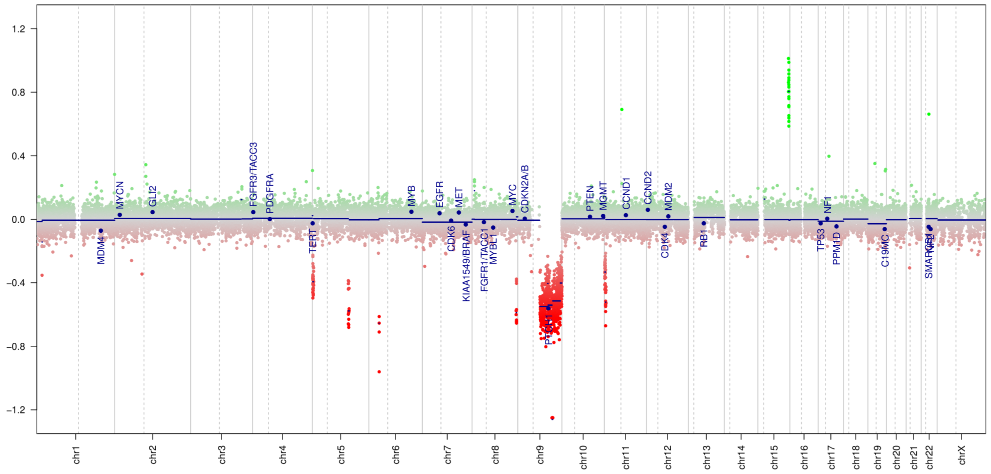


LFS2

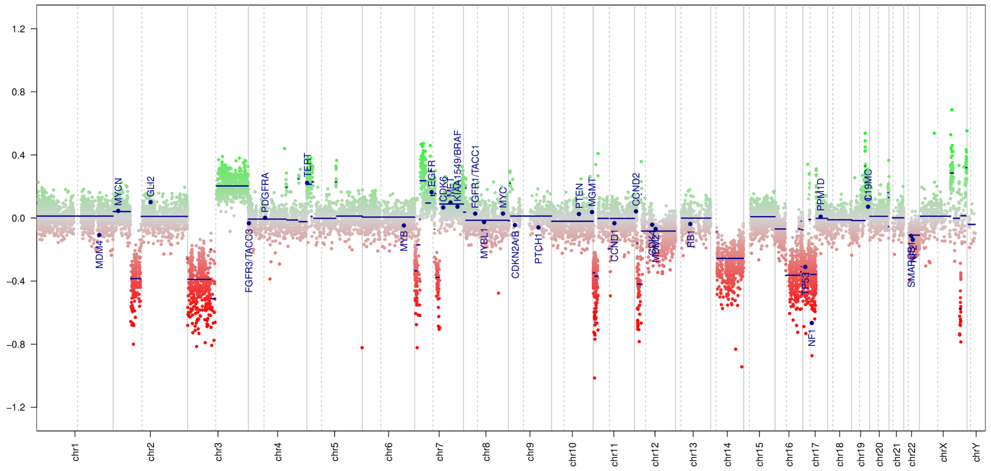


LFS3 and LFS8

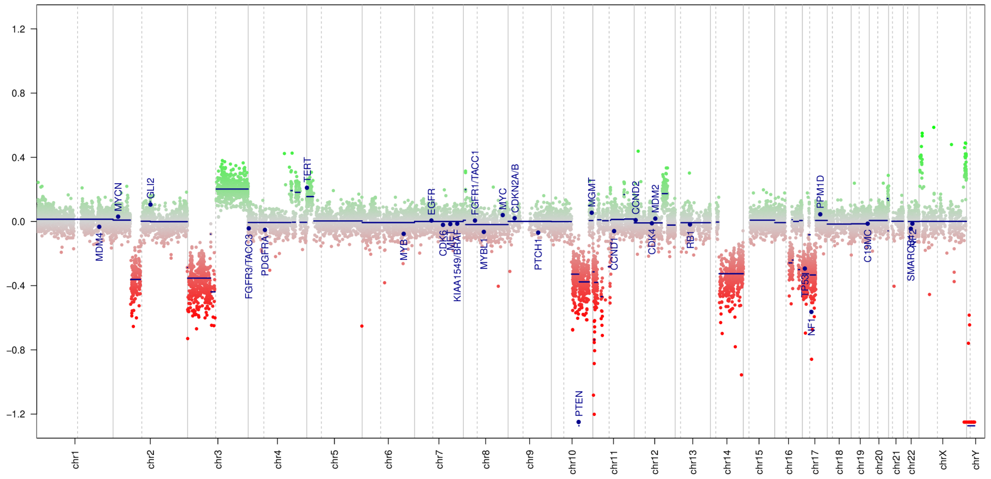


LFS4

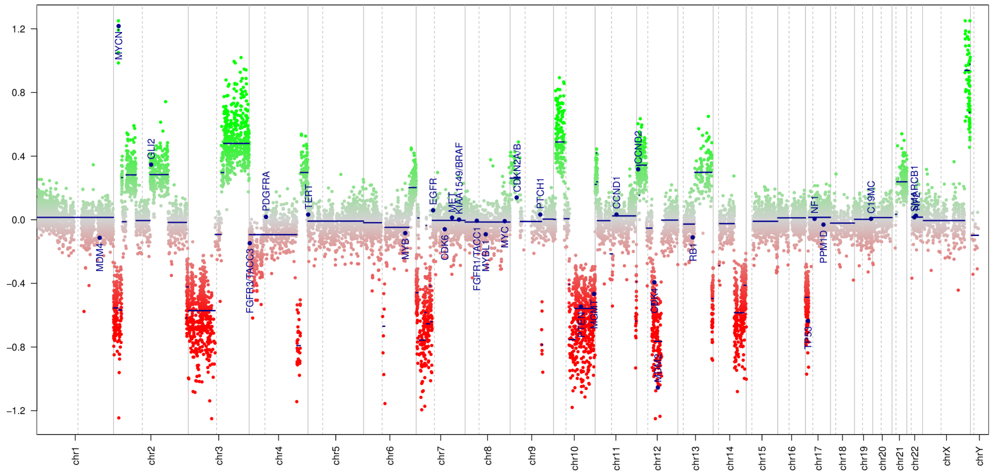

#### LFS5

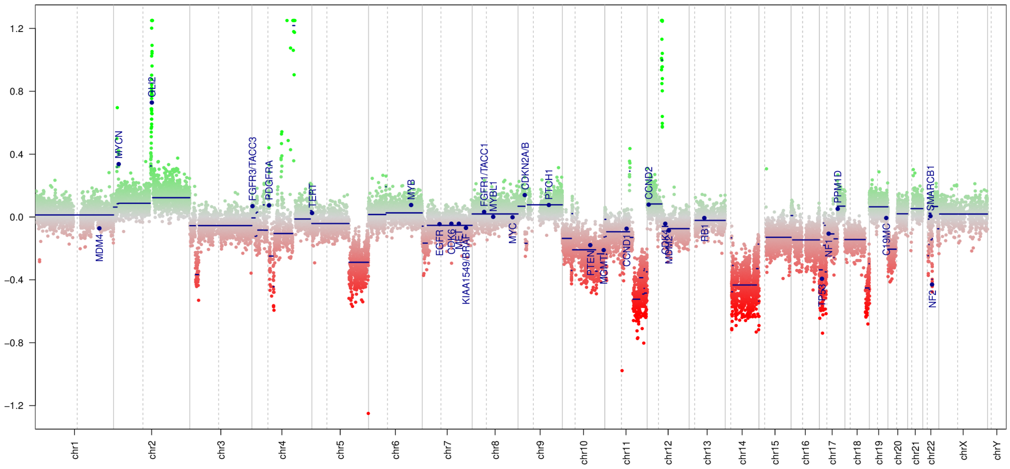


#### LFS6 and LFS7

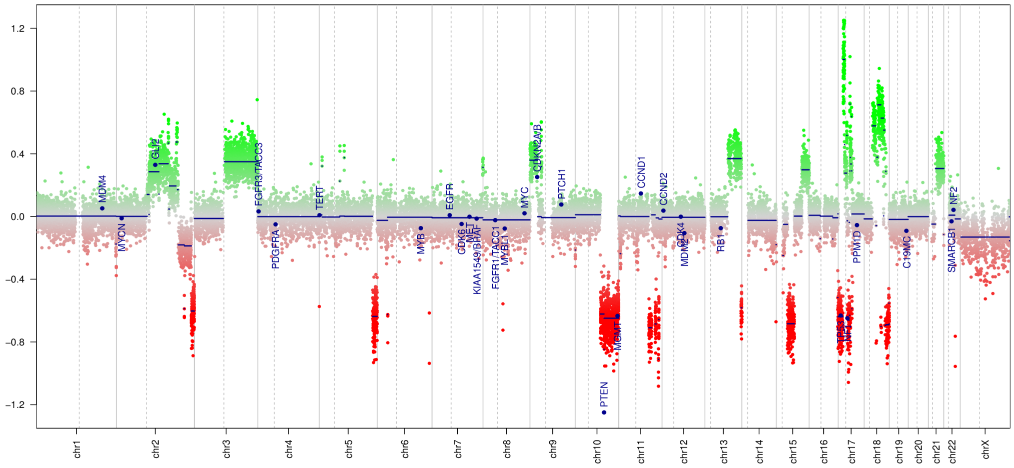


# S1

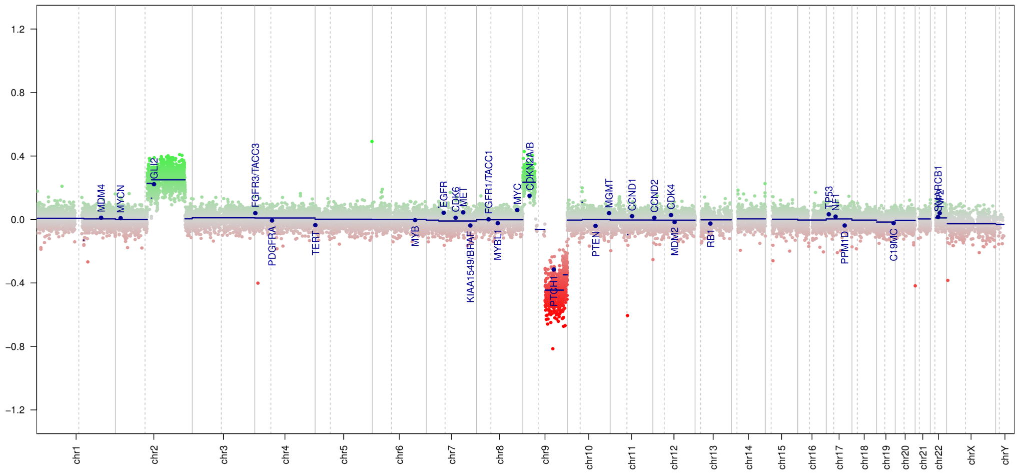

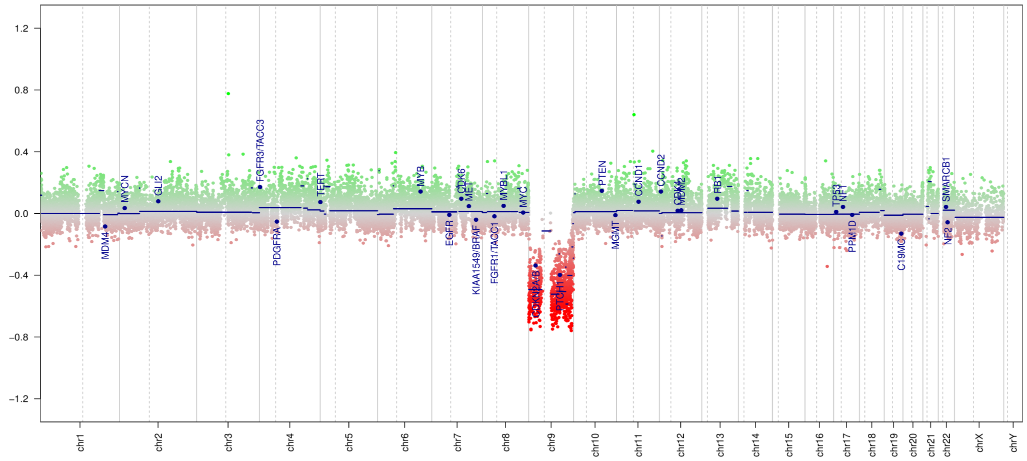

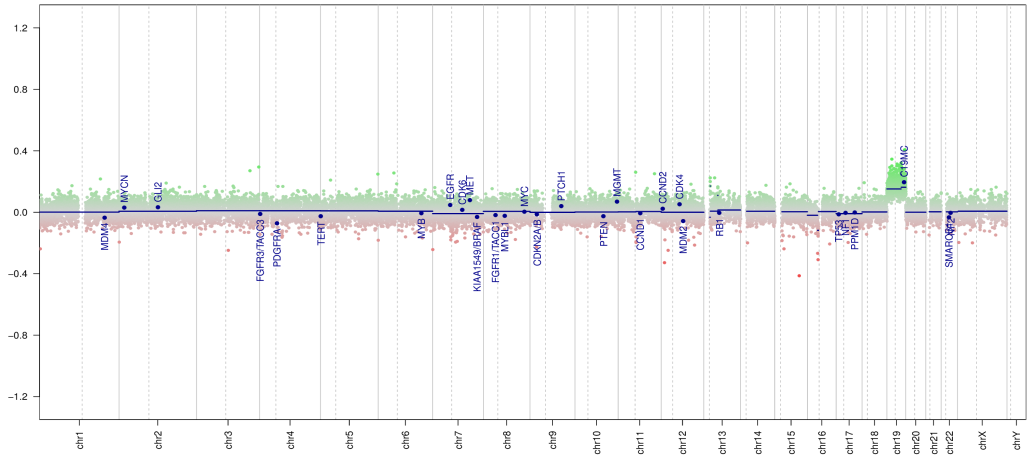

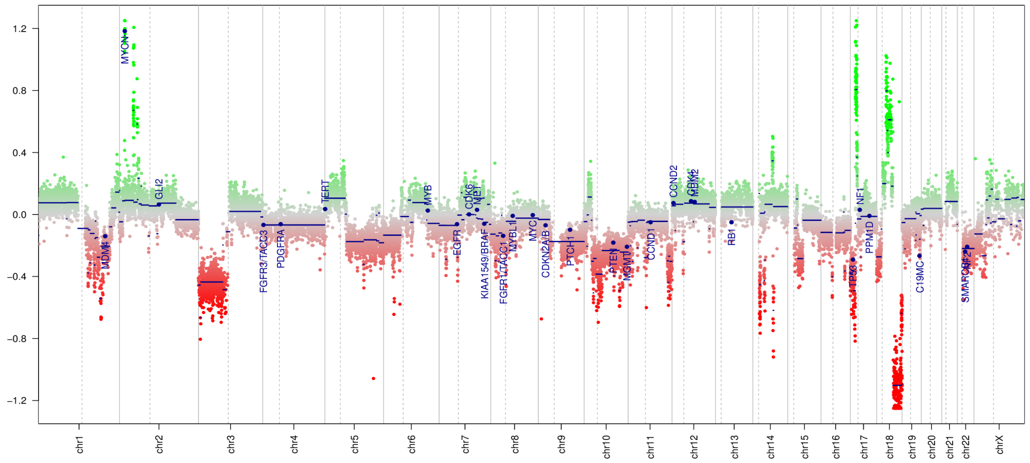

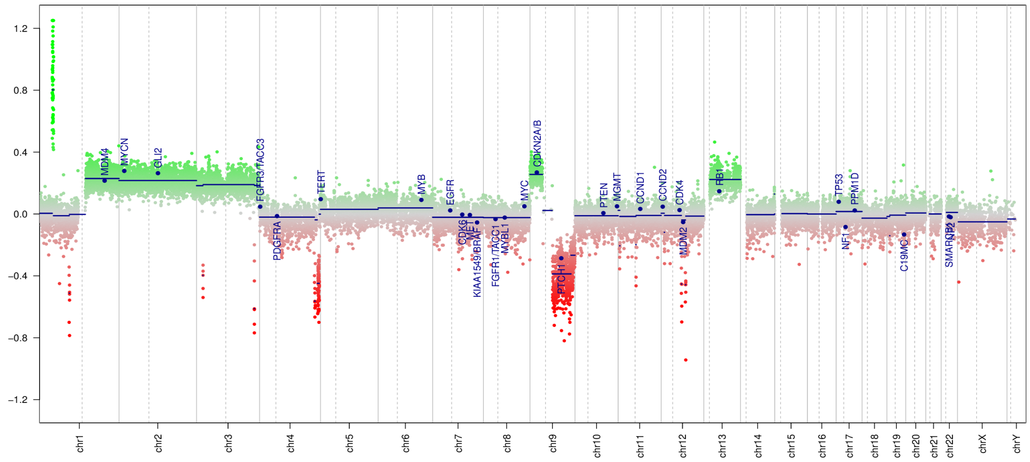

### Figure S2

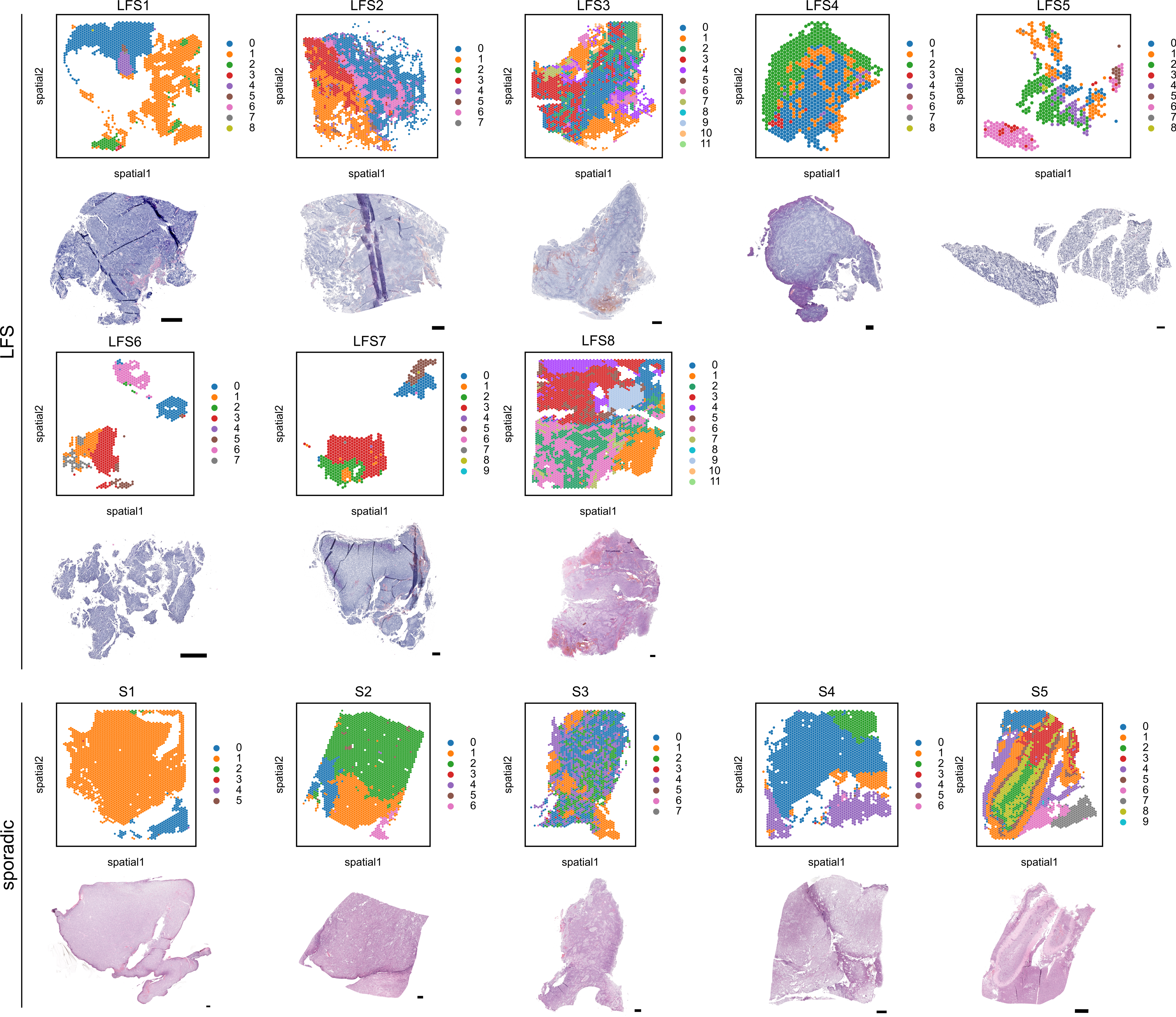

### Figure S3

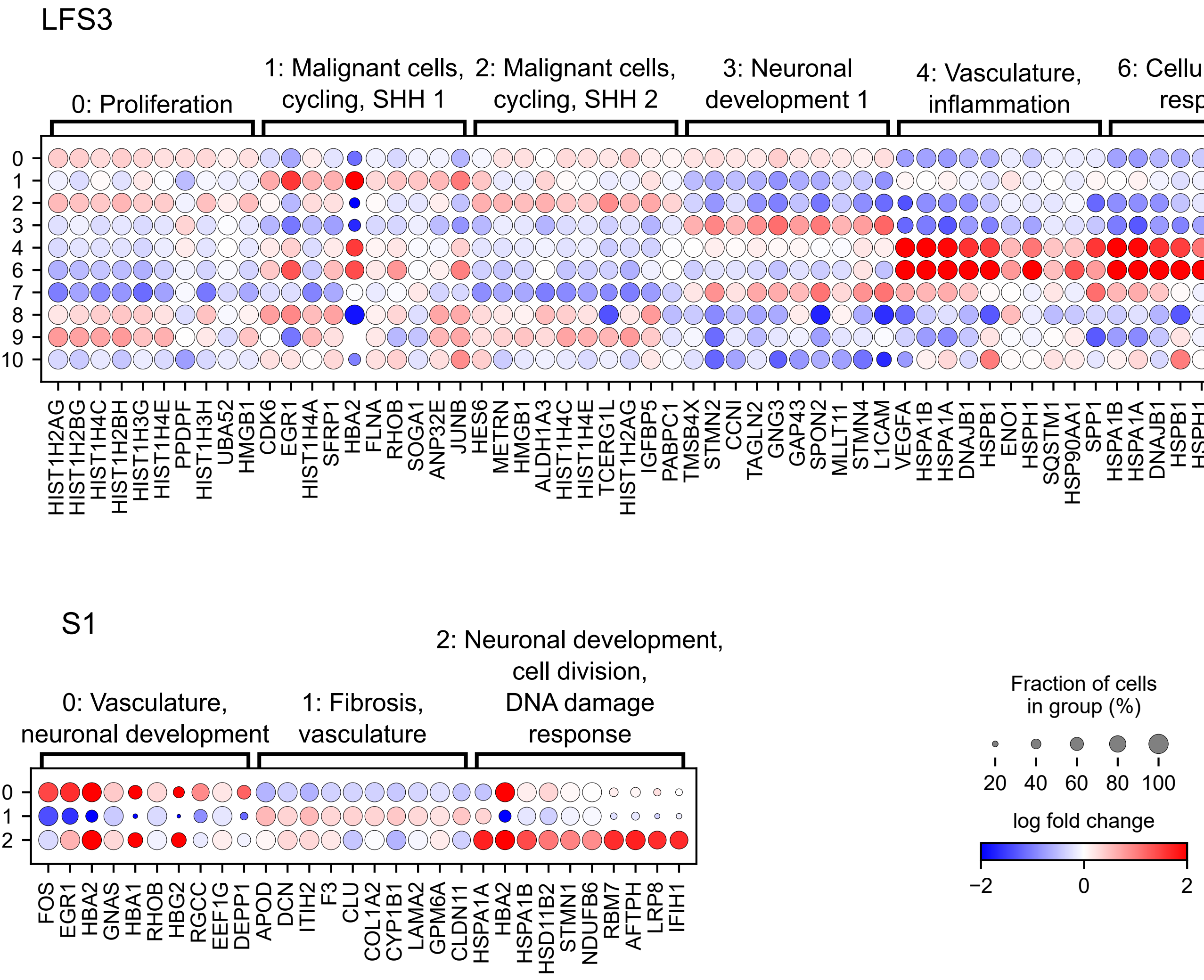

### Figure S4

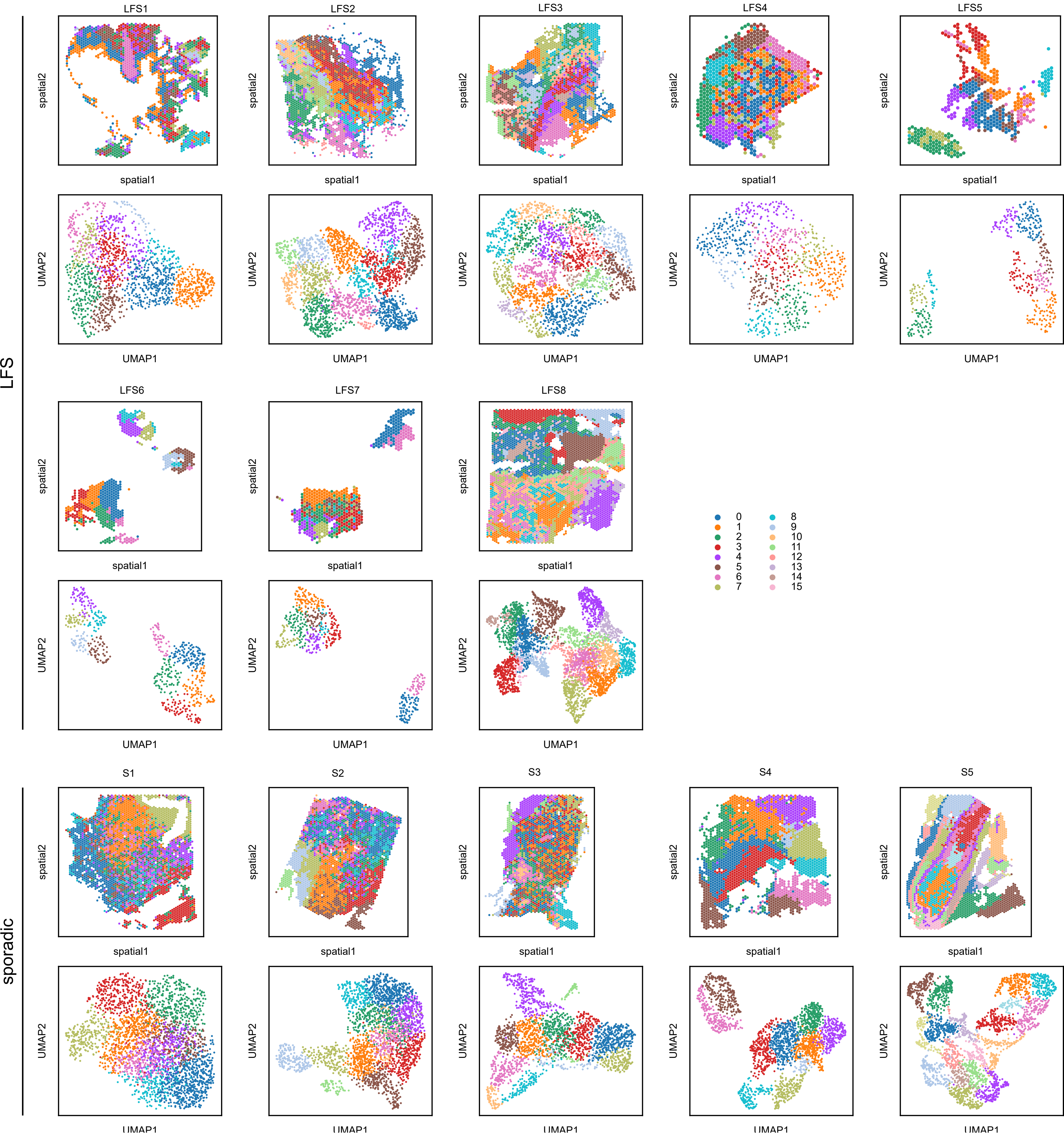

### Figure S5

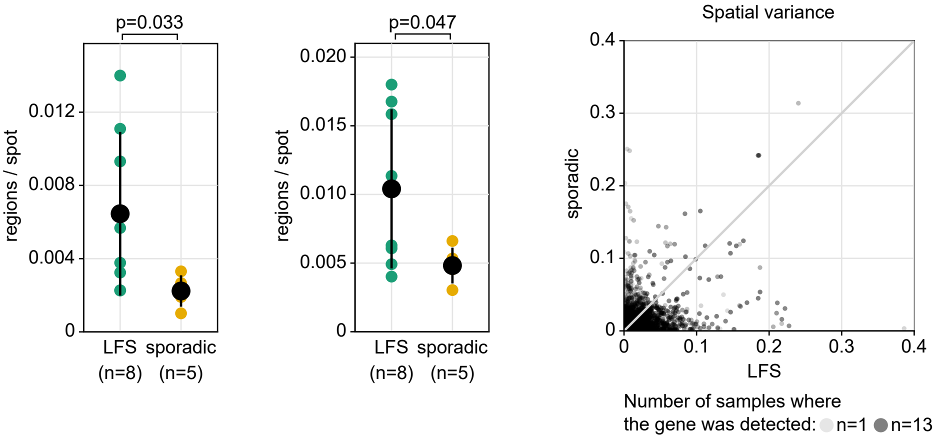

### Figure S6

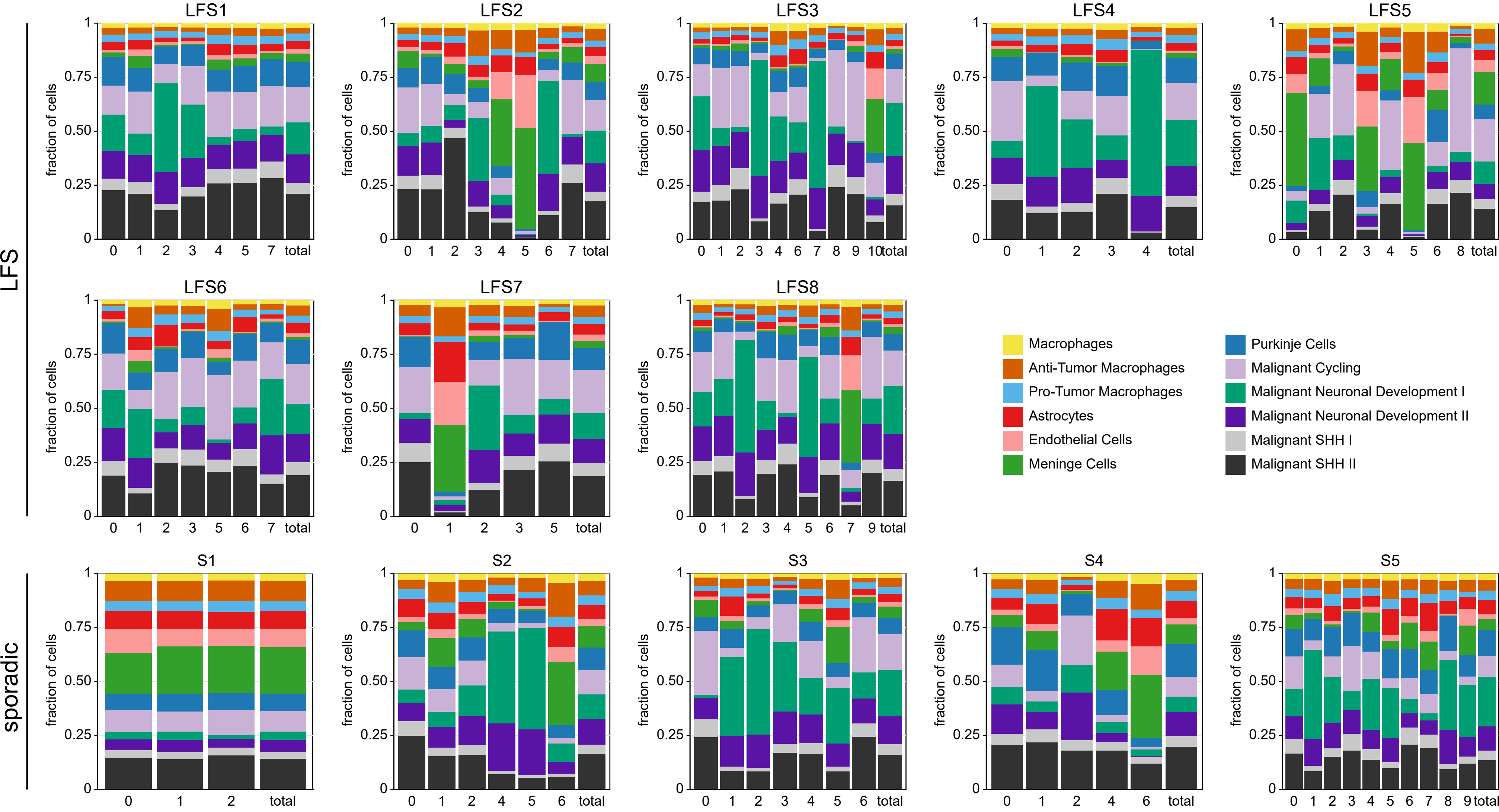

### Figure S7

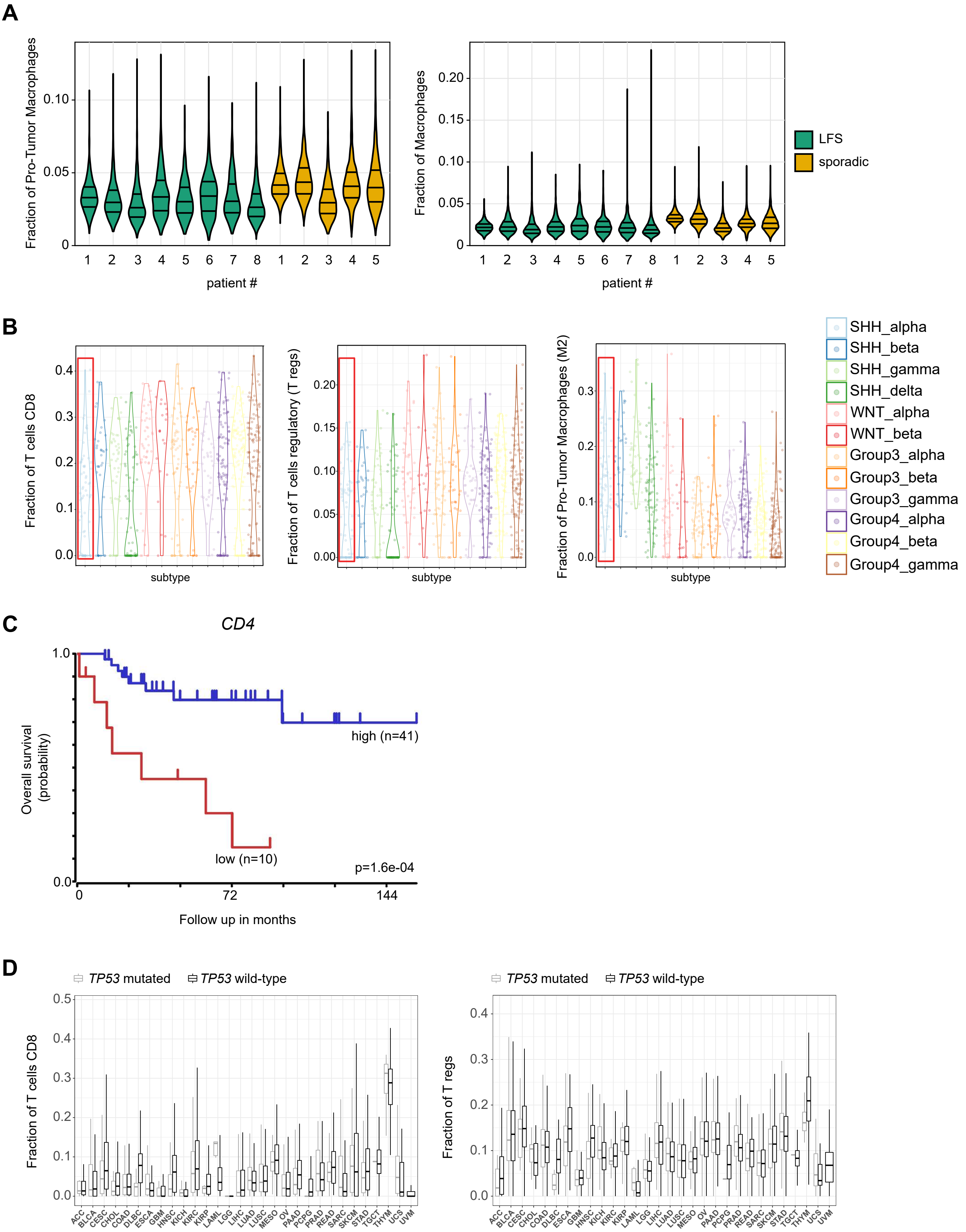

### Figure S8

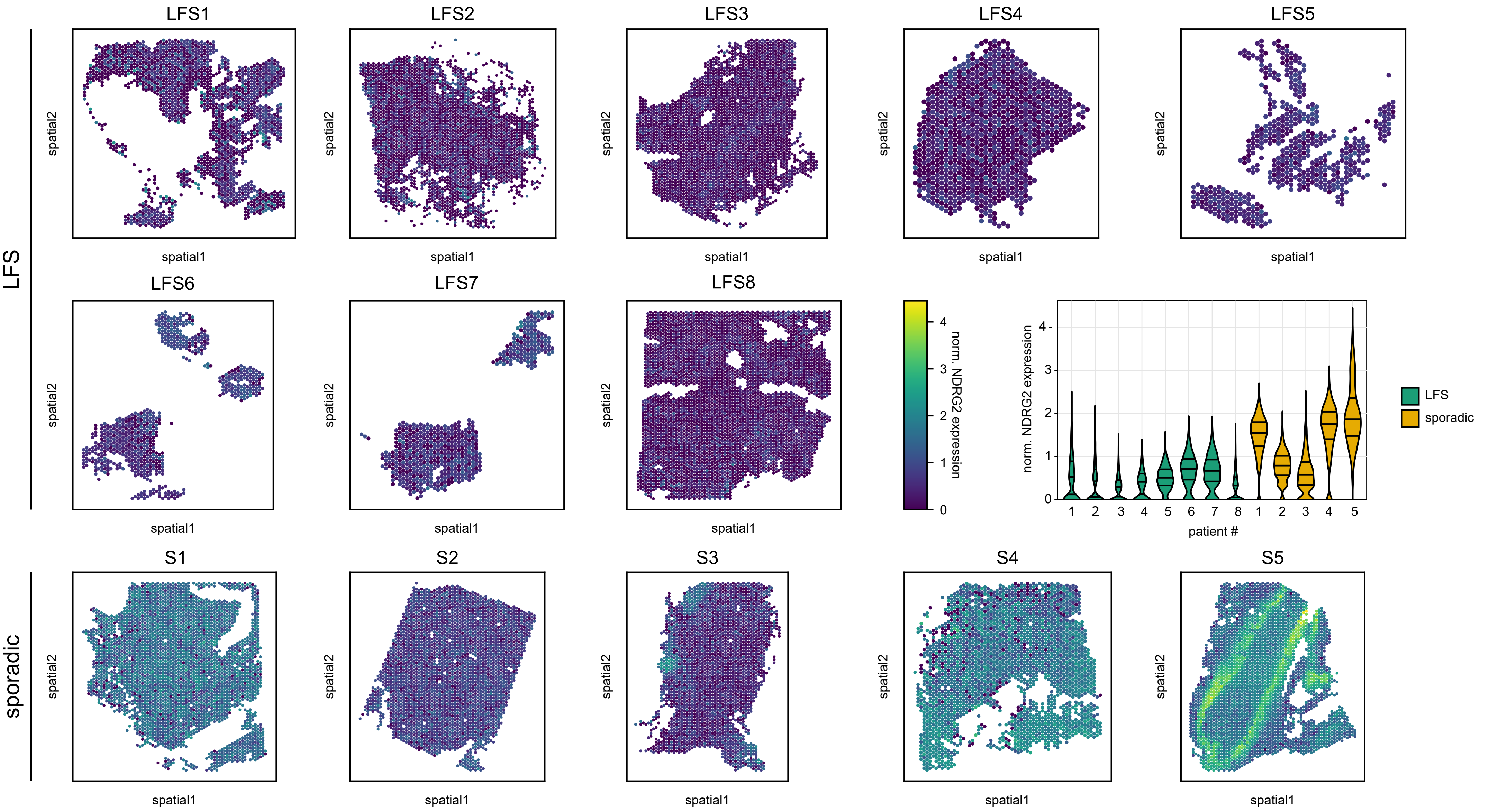

### Figure S9

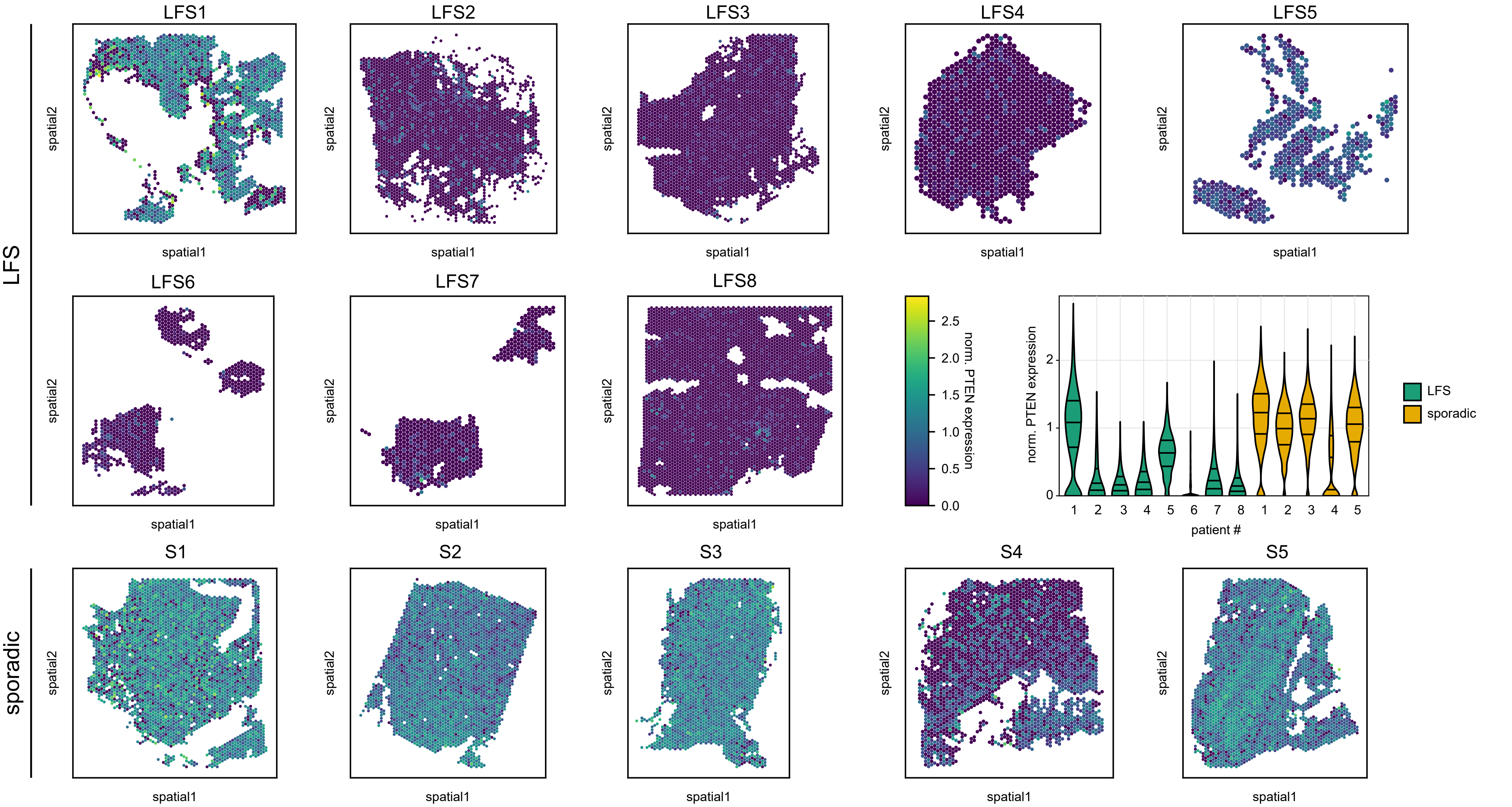

### Figure S10

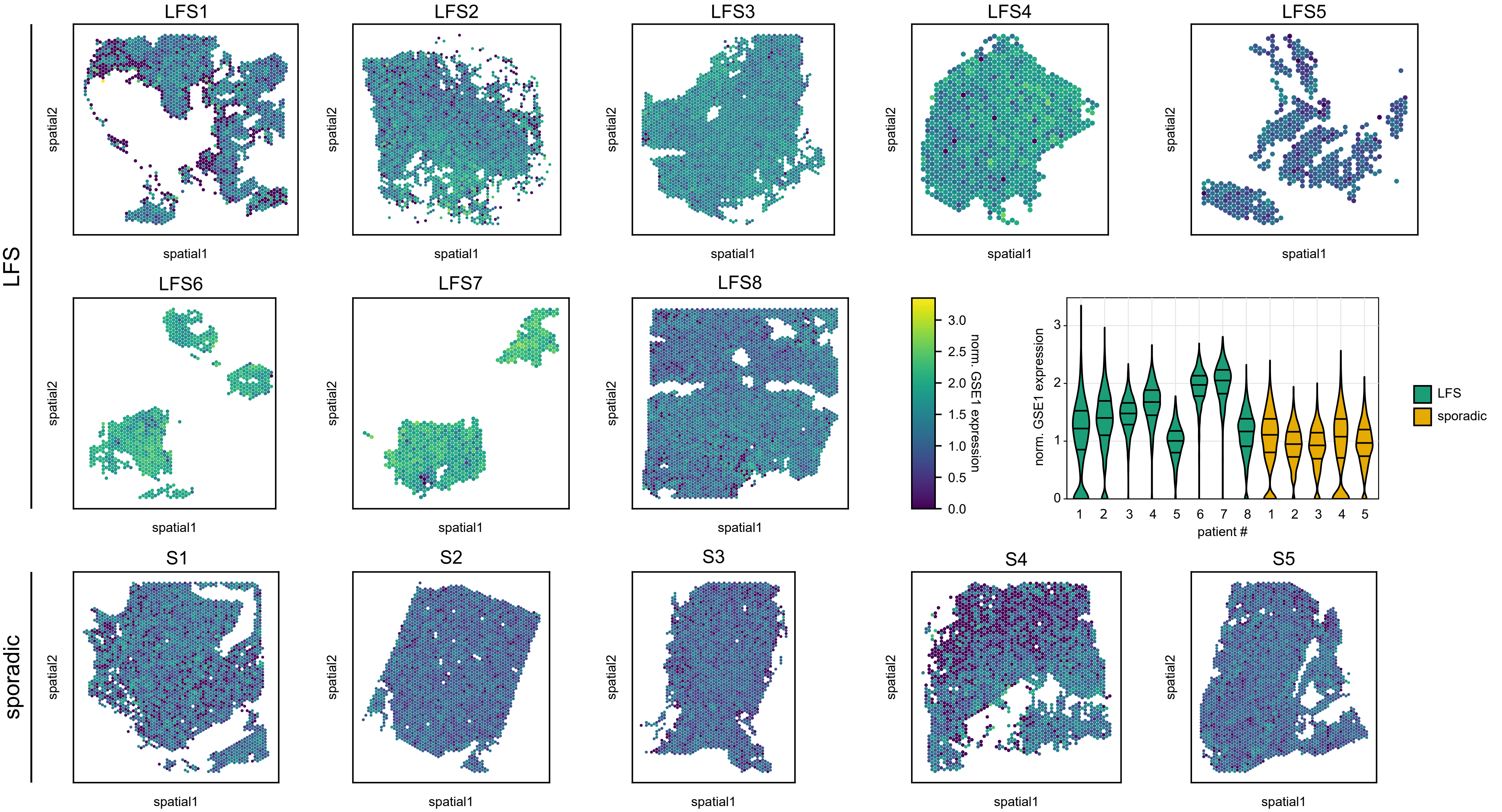

### Figure S11

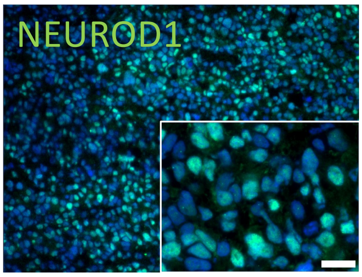

### Figure S12

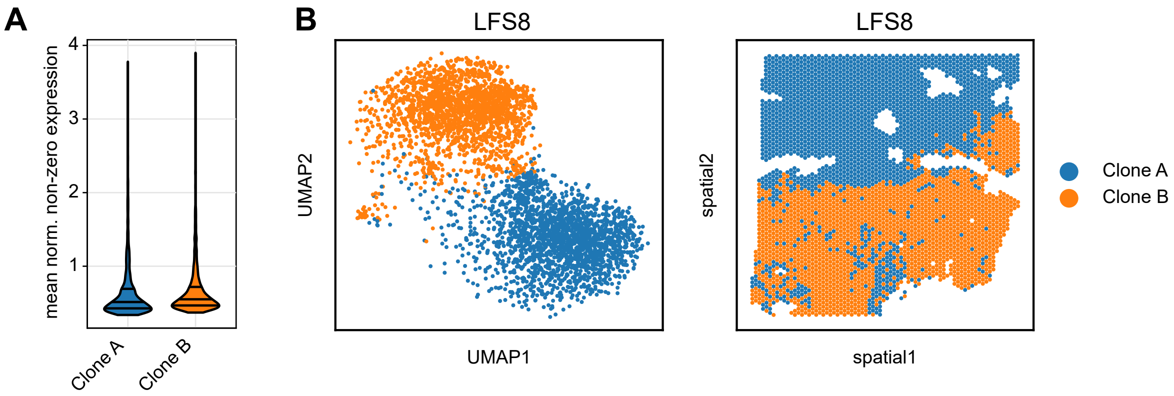

### Figure S13

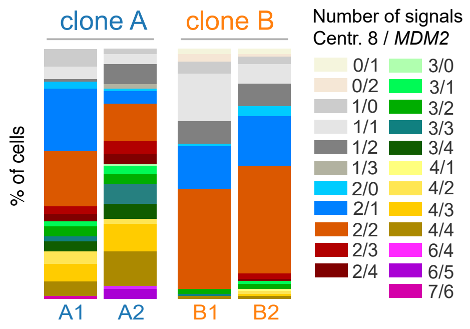

### Figure S14

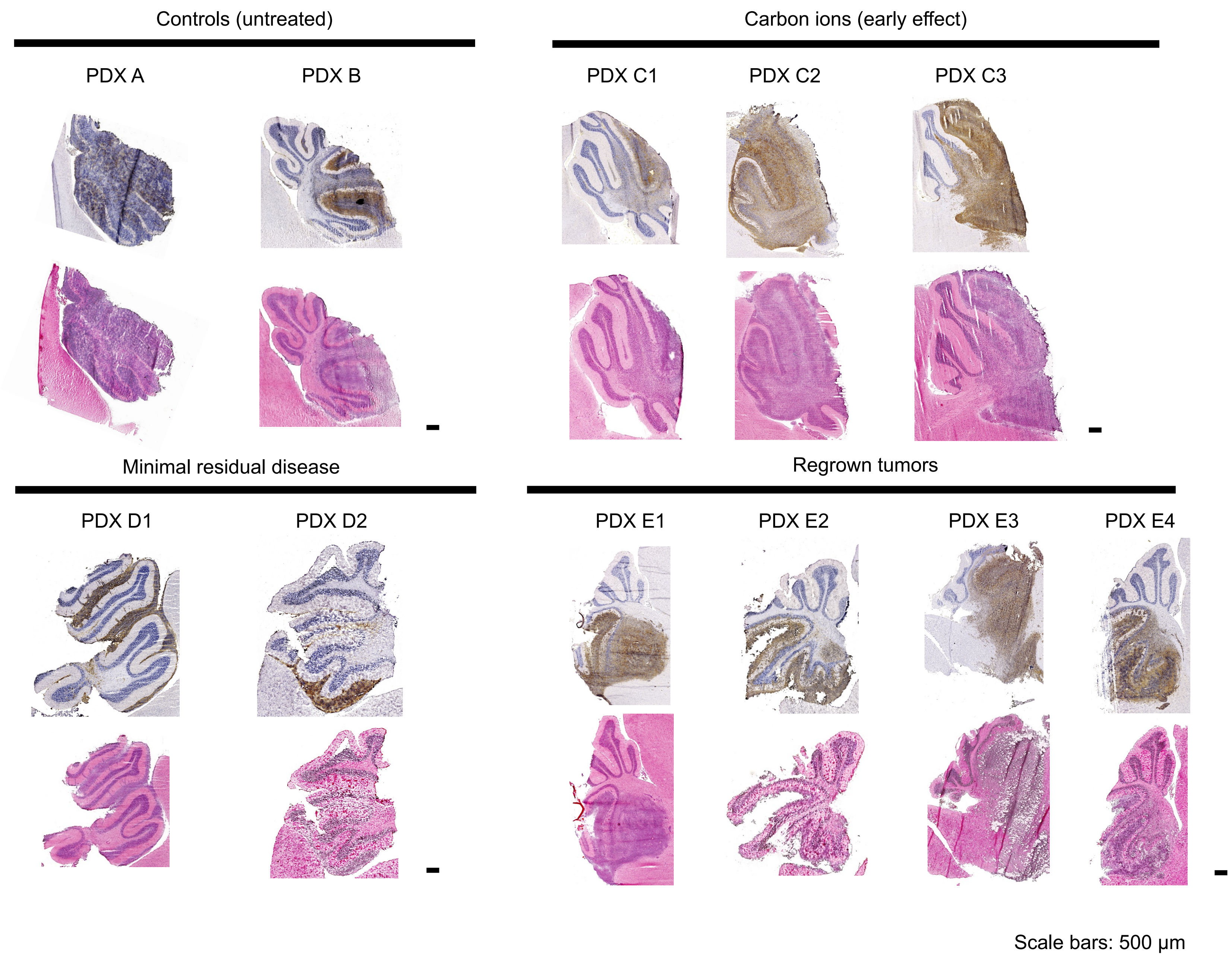

### Figure S15

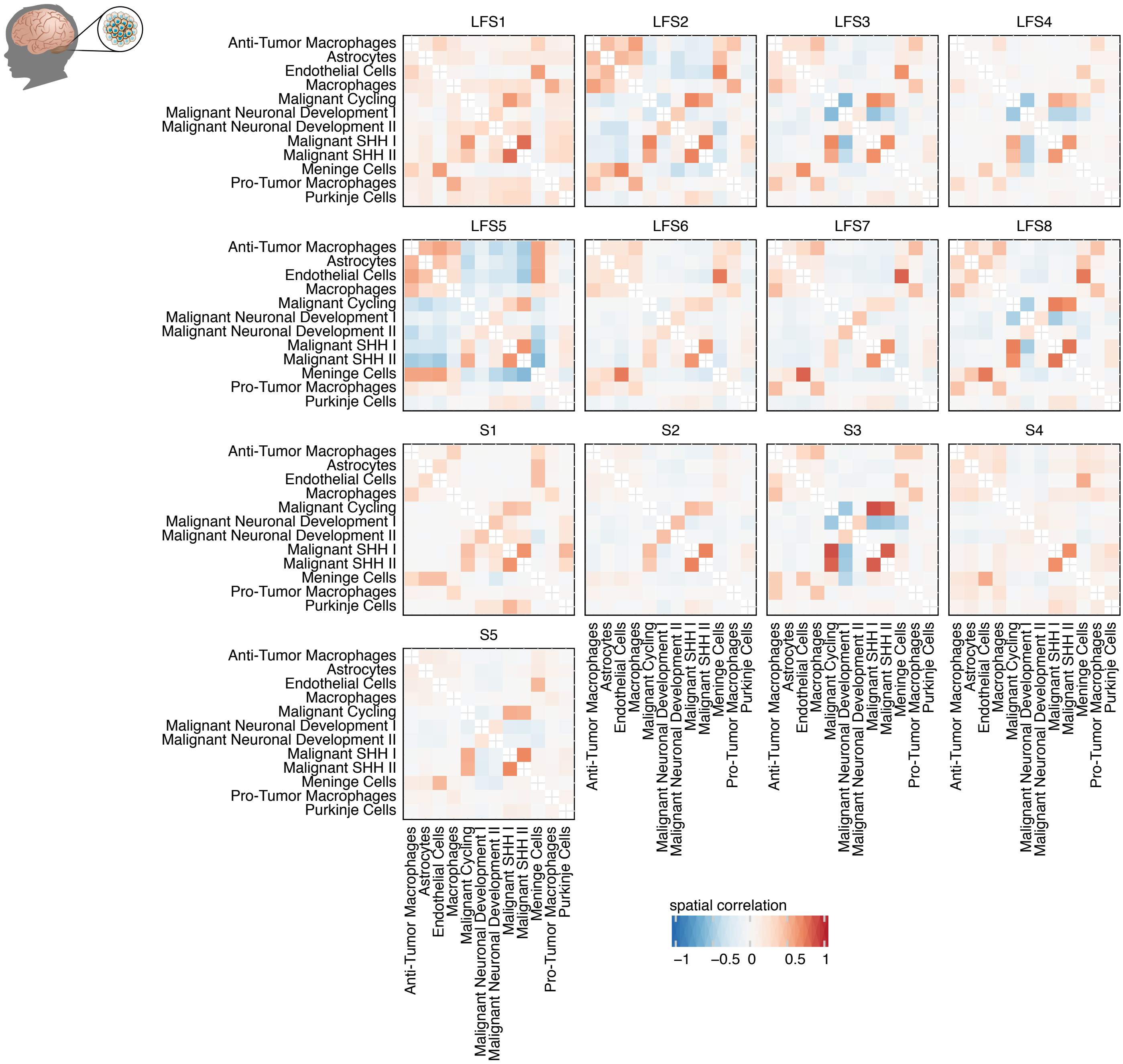

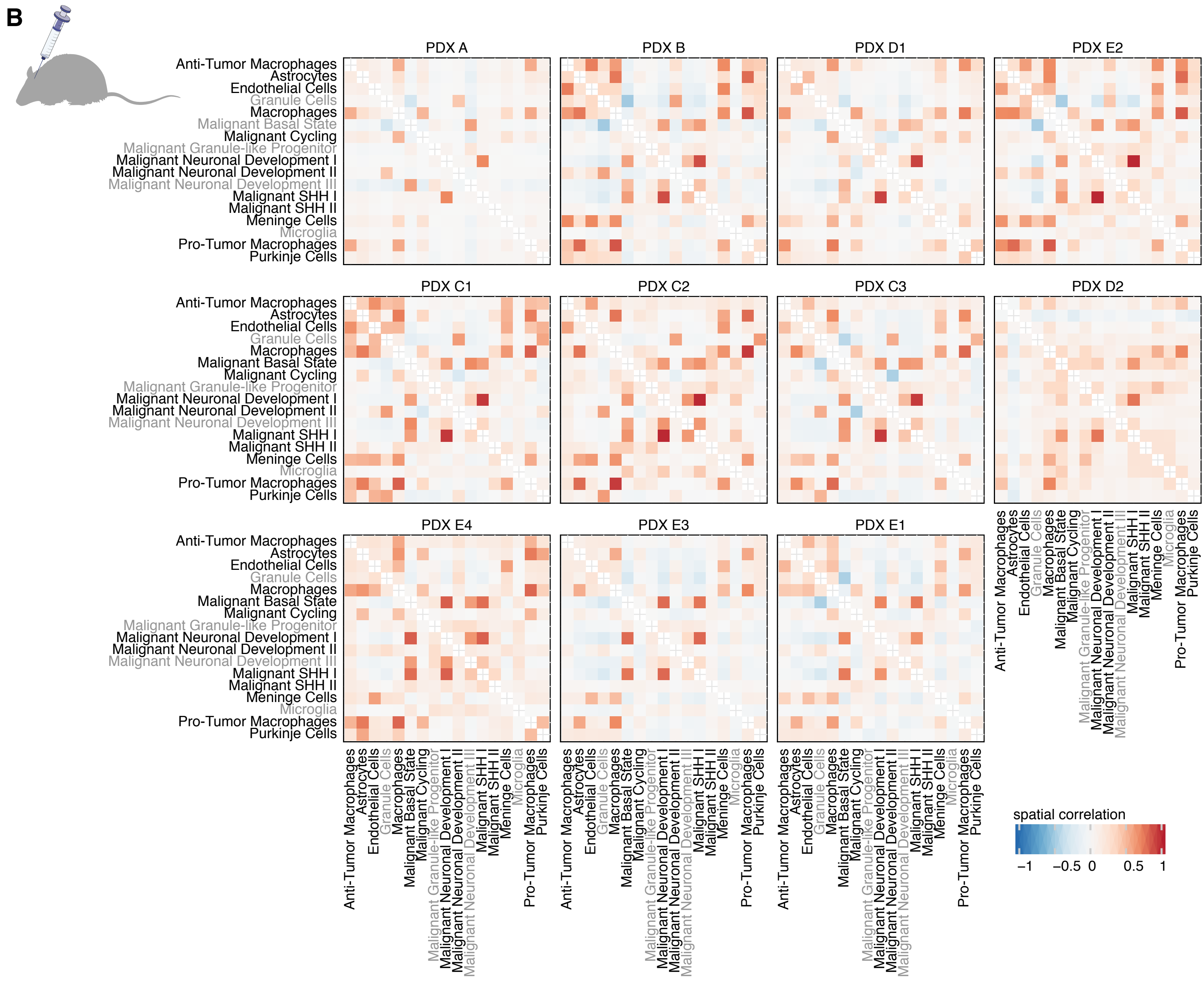

### Figure S16

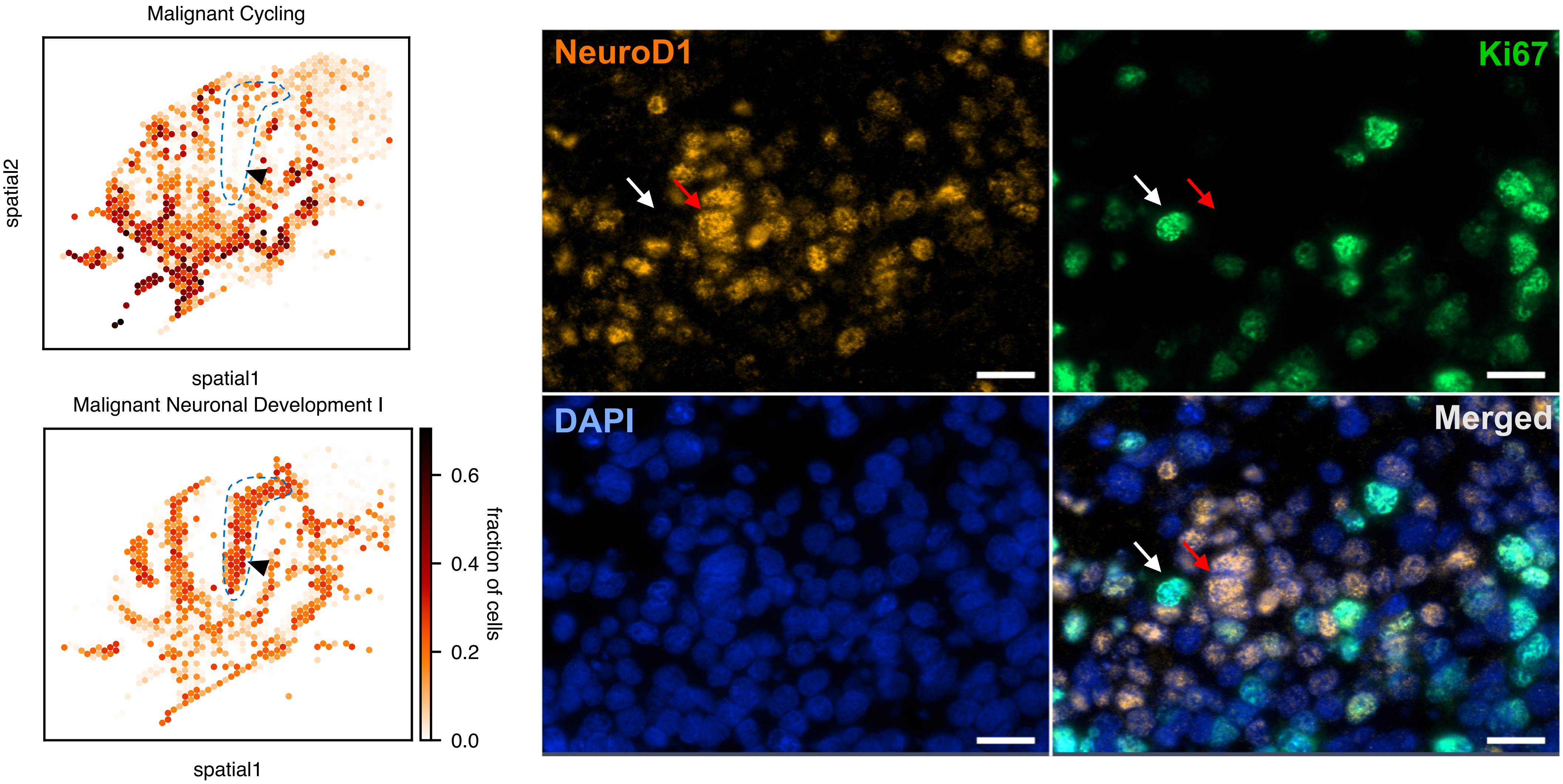

### Figure S17

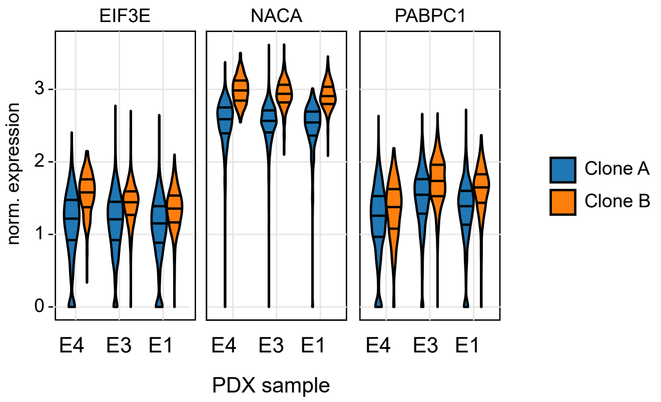

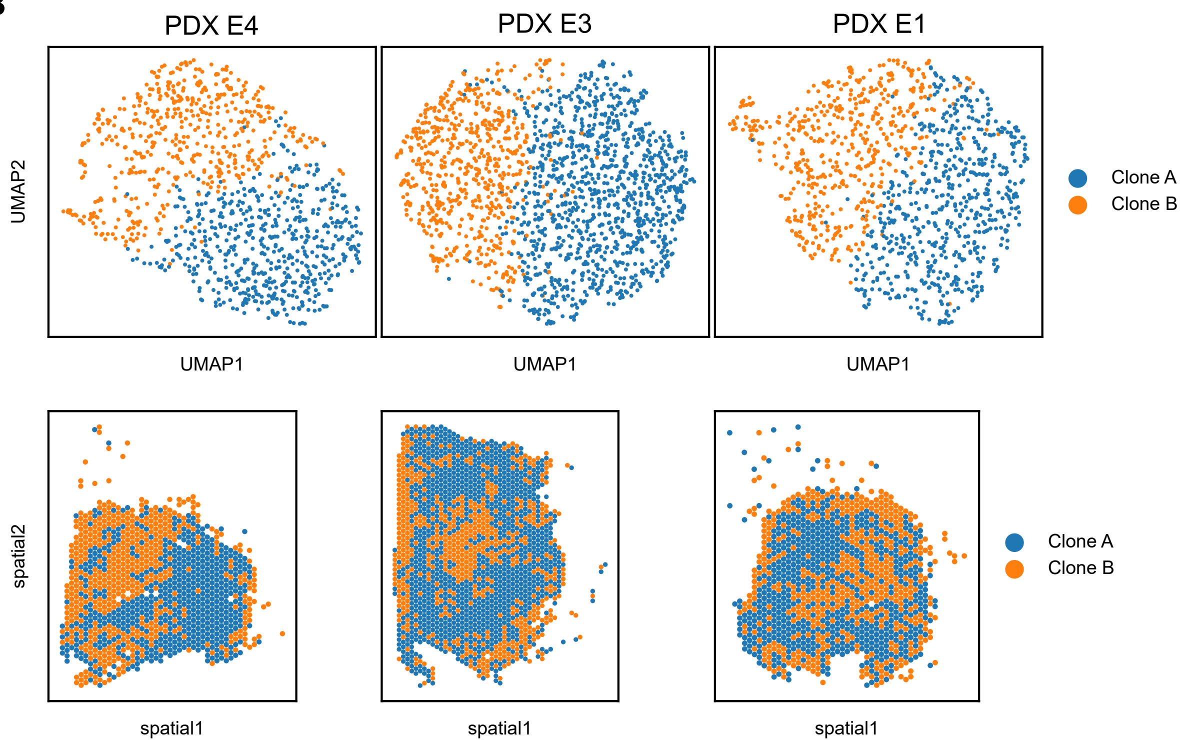

### Figure S18

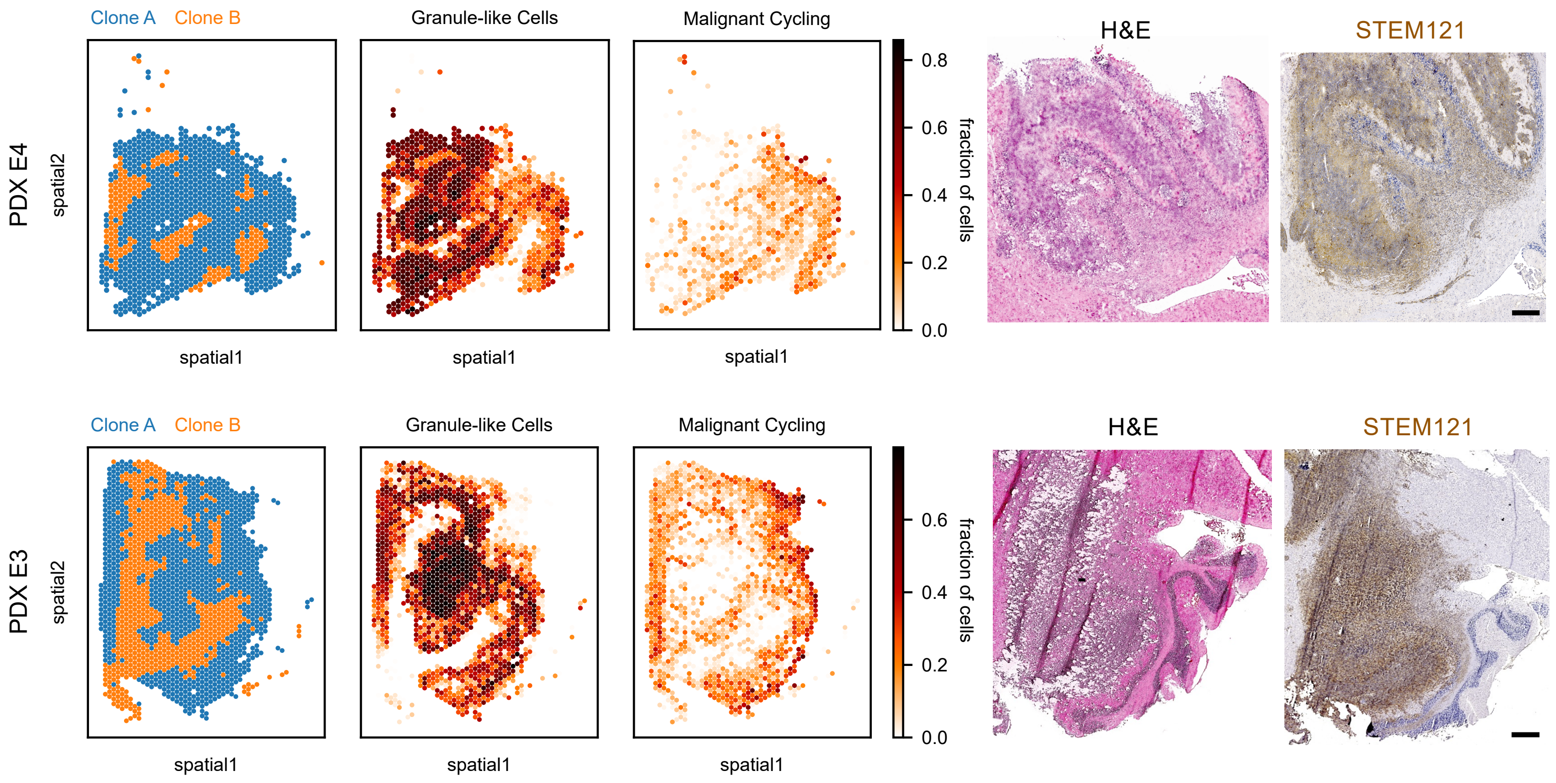

### Figure S19

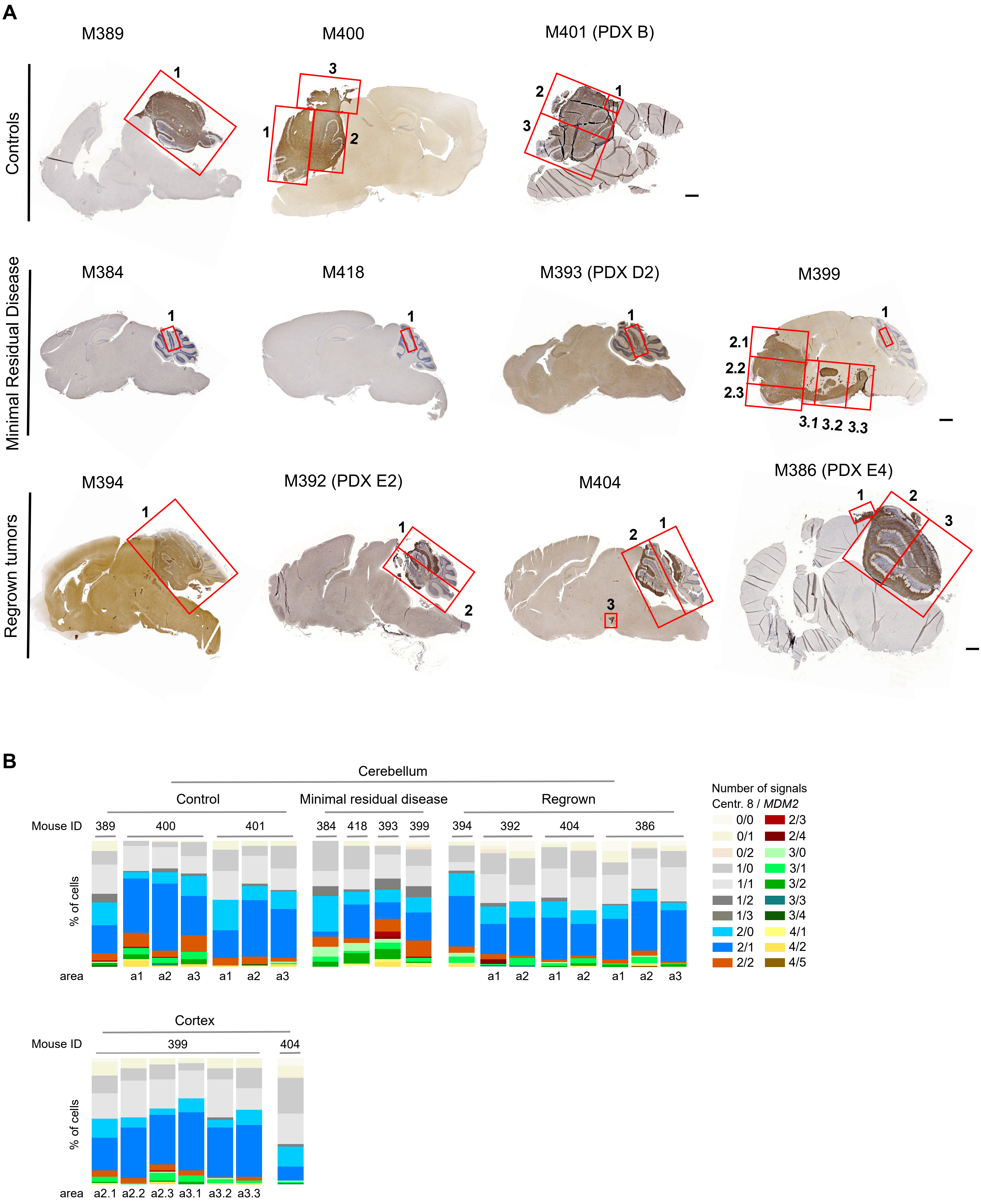

### Figure S20

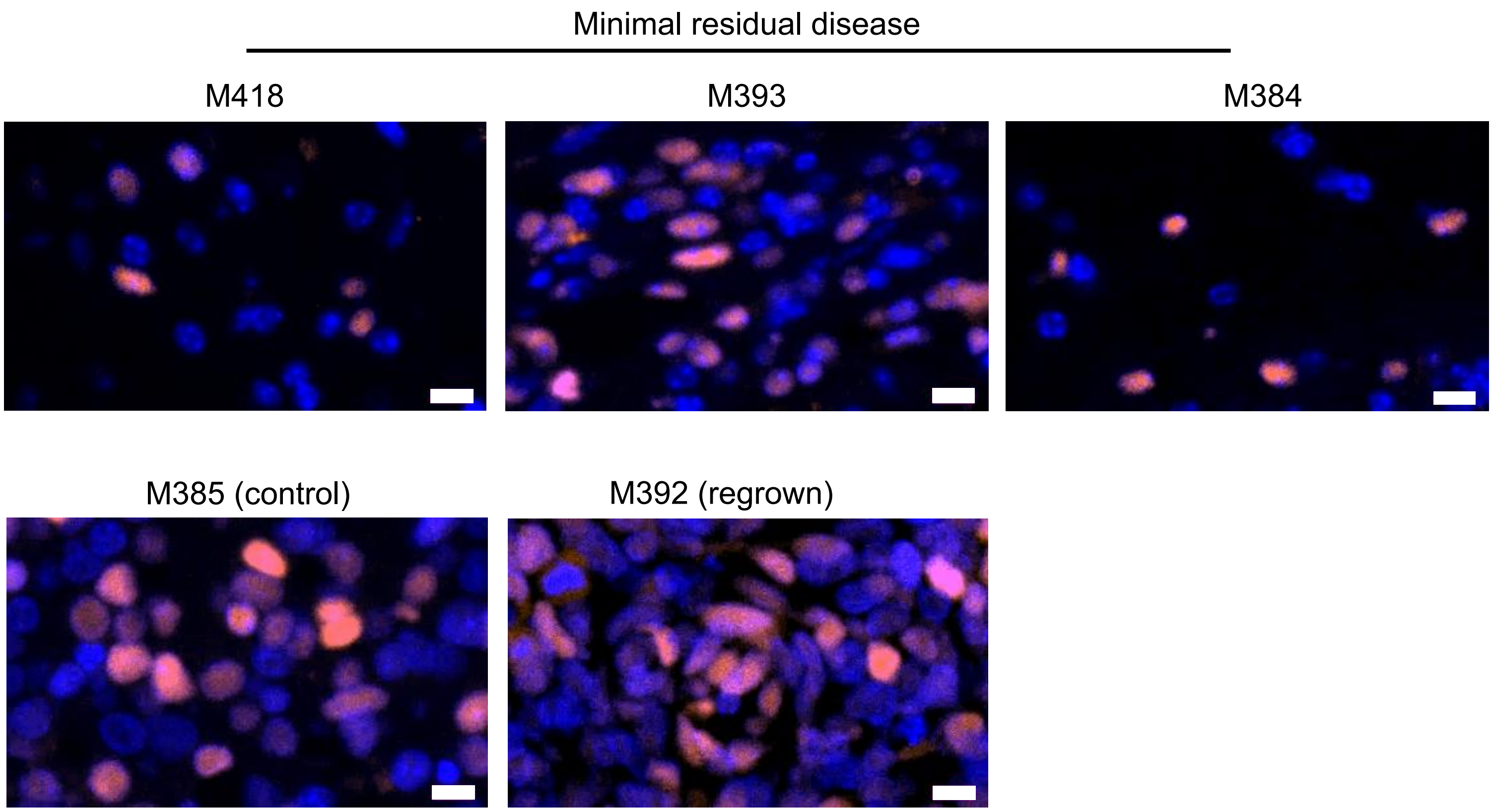

### Figure S21

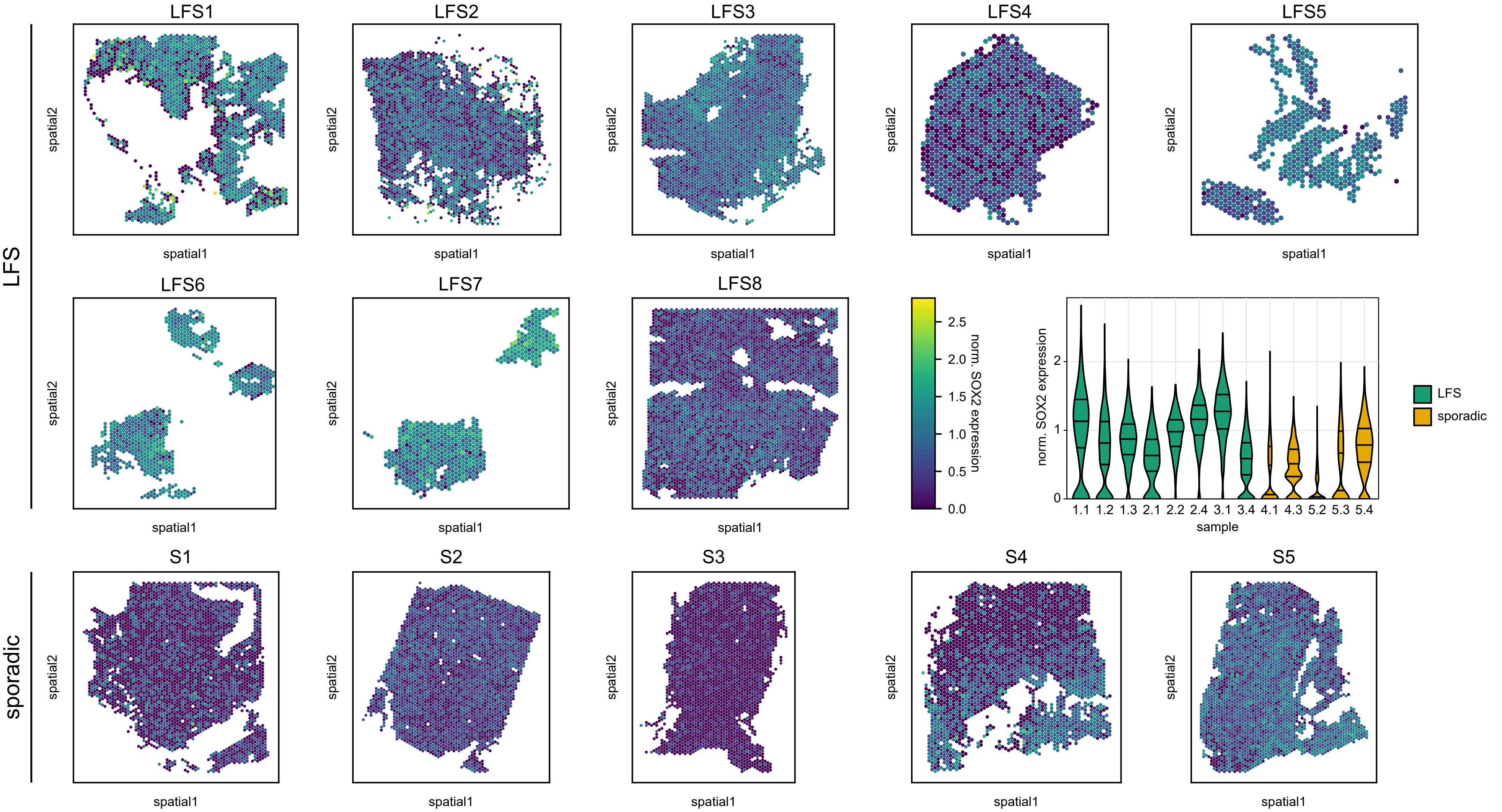

### Figure S22

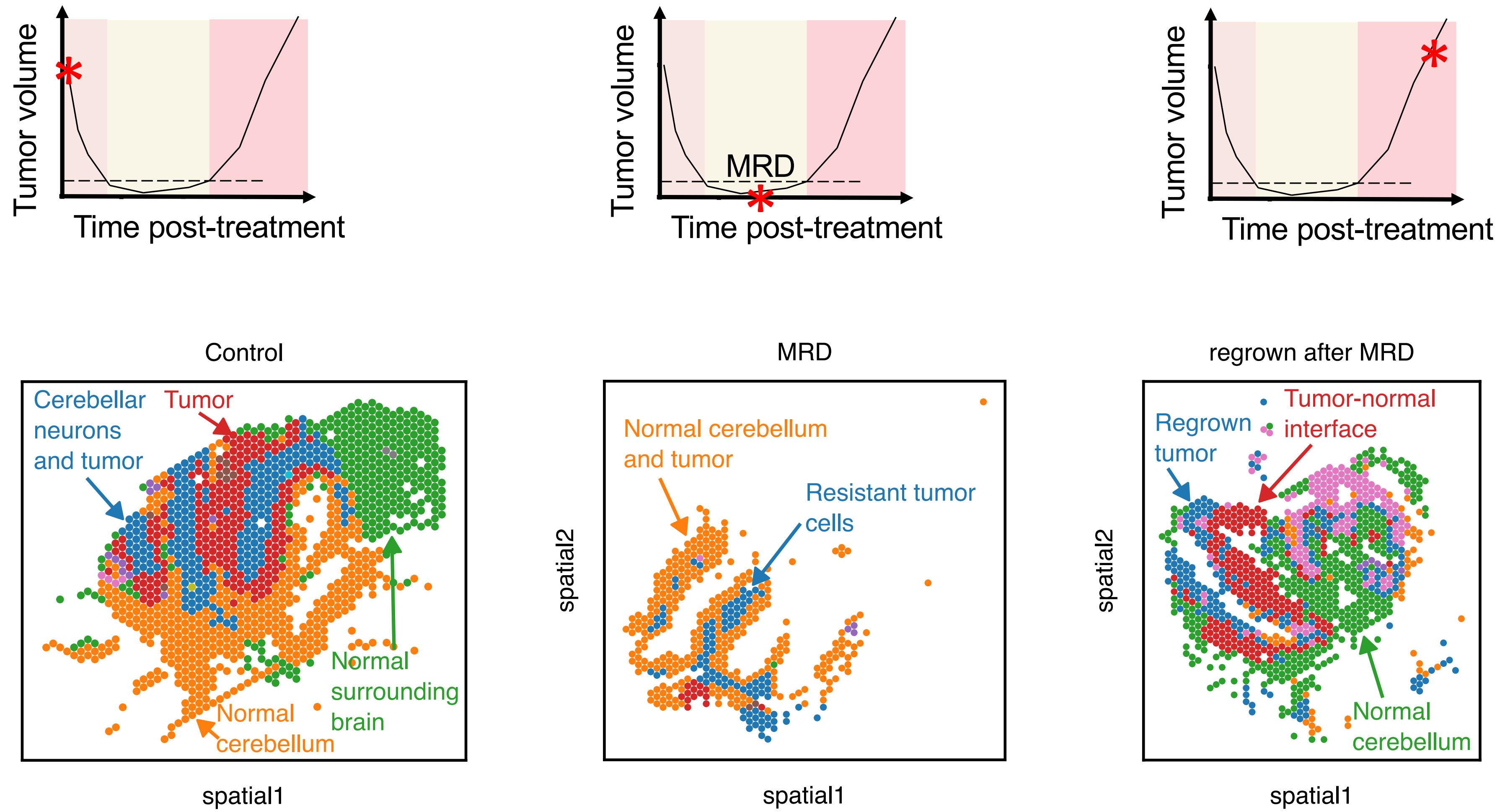

### Figure S23

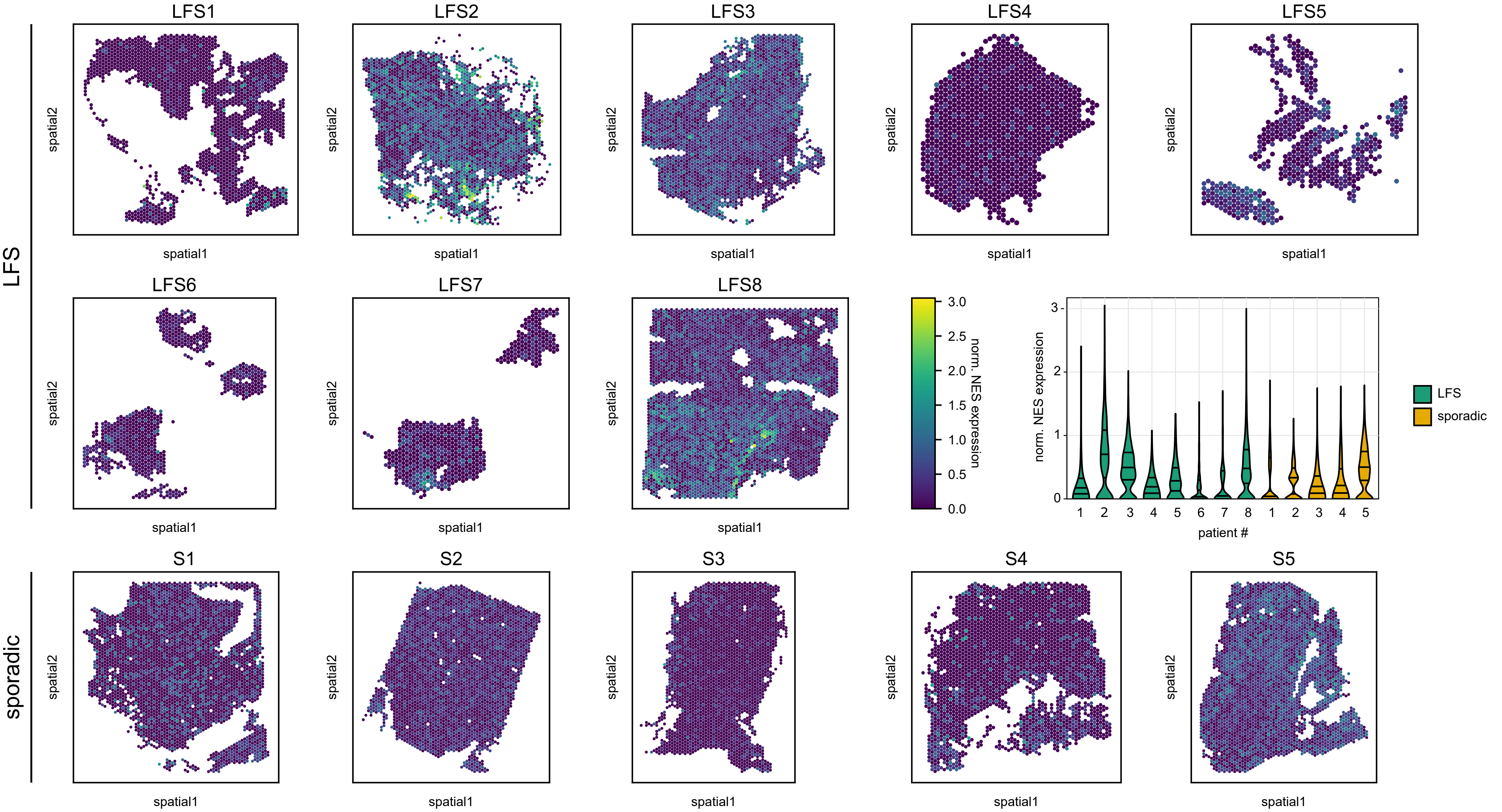

### Figure S24

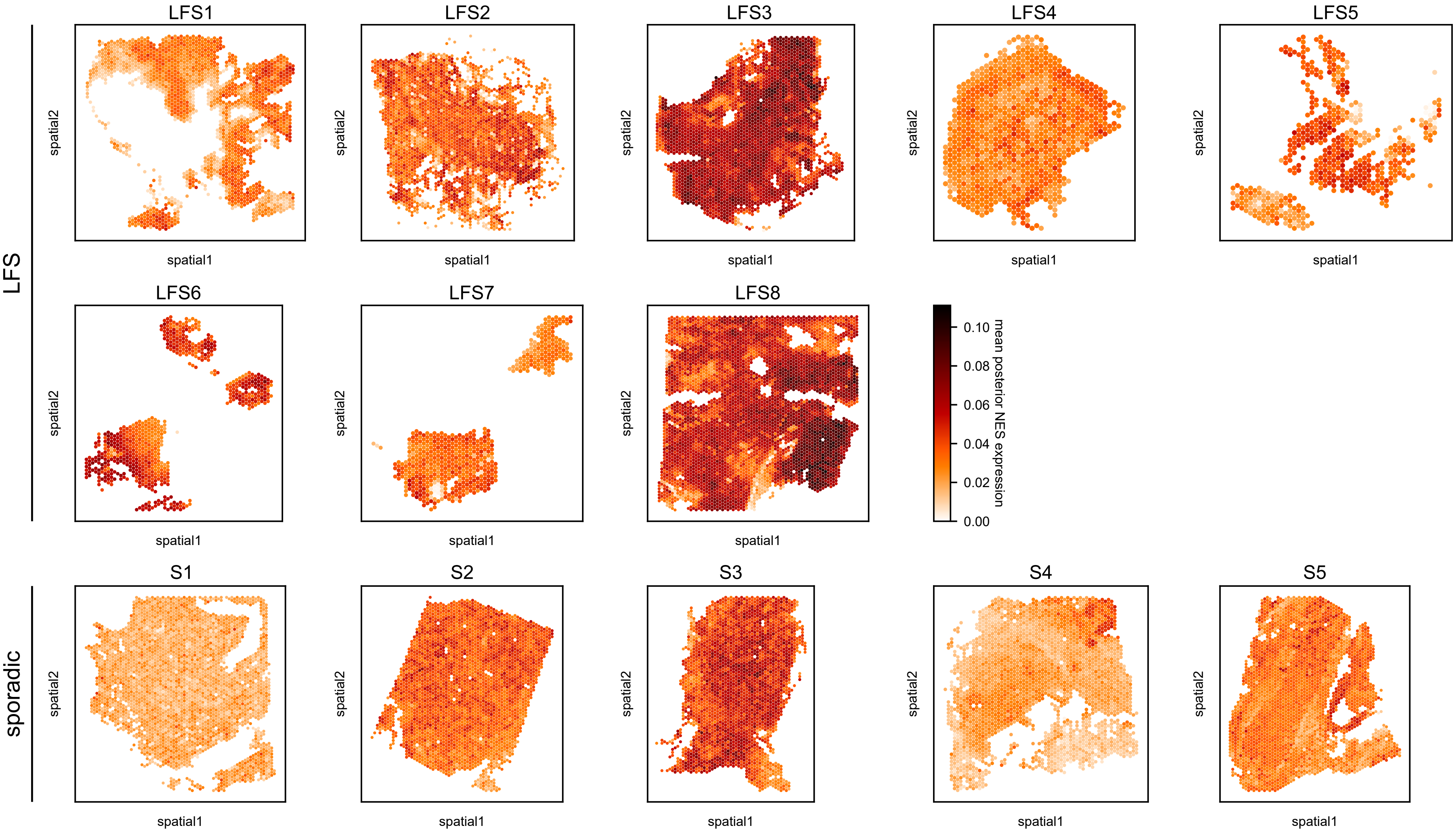
